# Translational Proteomics for Transfusion Medicine: Resolution of the IVIG Proteomes of Different Geographically Sourced and Prepared IVIG Immunotherapies

**DOI:** 10.1101/2021.04.21.440739

**Authors:** Garry W. Lynch, Anna M. Fitzgerald, Bradley J. Walsh, Natalie Kapitza, John S. Sullivan

**Affiliations:** The School of Medical Sciences, Faculty of Medicine and Health, of The University of Sydney, Camperdown, NSW, 2006, Australia; The School of Veterinary Science, Faculty of Science, of The University of Sydney, Camperdown, NSW, 2006, Australia; The Marie Bashir Institute for Infectious Diseases, Immunity and Biosecurity, of The University of Sydney, Camperdown, NSW, 2006, Australia; Minomic International Ltd, Macquarie Park, NSW, 2113, Australia; Whitlam Orthopaedic Research Centre, Ingham Institute for Applied Medical Research, Liverpool, NSW, 2170, Australia; Sydney Medical School, Faculty of Medicine and Health, The University of Sydney, Camperdown, NSW, 2006, Australia; School of Medical Science, University of New South Wales, Randwick, NSW, 2052, Australia

**Keywords:** IVIG, IGIV, Cytokine, Immunoglobulin, Antibody, Transfusion Medicine, Proteomics, Multiplex, Mass Spectrometry

## Abstract

Intravenous Immunoglobulin’s (IVIG’s) are prepared from thousands of donor plasmas and as a result comprise an extreme broad mix and depth of Antibody (Ab) specificities. IVIG formulations available in Australia are from both local and overseas donor sources and extracted using a variety of purification methods and immunoglobulin purities, with the Australia-derived and prepared IVIG listed at >98% and the overseas-derived preparations at ≥ 95%. Because of these differences it was predicted that the formulations might individually vary in composition and that together with obvious genetic and geographic antigenic (Ag) environment differences may result in notable variability between formulations. Hence a focussed comparative proteomic profiling of IVIG formulations was undertaken to identify notable similarities and differences across products. Comparisons between formulations did reveal marked qualitative differences in 2D-gel Antibody profiling that included parameters of isoelectric charge (pI), as well as immunoglobulin (Ig) monomer to dimer ratio variability between products, including high molecular weight (MW) immunoglobulin multimers for some. These notable differences were in part quite likely a product of the respective purification methods used, and capacity to select (or de-select) for antibodies of such different properties. Furthermore, for identification of non-Ig proteins carried over from plasma through purifications Mass spectrometry was performed. This identified a few such ancillary proteins, and their identities, in general, differed between formulations. Proteins detected included the most abundant protein of plasma, albumin, as well as other mostly large and abundant proteins; RAG1 - V(D)J recombination activating protein1, gelsolin, complement Factor-B, serotransferrin, tetranectin, NADH ubiquinone oxidoreductase, caspase 3 and VEGFR1. An alternate strategy used commercial Multiplex xMAP assay to detect cytokines, which are small and present in plasma at trace but highly active quantities. This revealed various different cytokine profiles across the formulations studied. The identification of additional proteins, and especially cytokines in IVIGs, is particularly notable, and the positive, negative or null biological relevance for clinical use, needs resolution. Collectively these findings reveal marked differences between Australian and overseas-derived (non-Australian) IVIGs in immunoglobulin composition and biochemical characteristics, and presence of additional carry-over proteins from plasma. These findings prompt the need for further evaluation of the micro-compositions of individual formulations. Perhaps detailed mining and improved comparative understanding of each IVIG formulation, may enable highly tailored and strategic clinical use of certain formulations that are personalised best-fit treatments for particular conditions. Such as in treatment of a neuropathy, as compared to another formulation, more suited for treating a particular infectious disease. The most salient and overarching study conclusion is need for caution in attributing equivalence across IVIGs.

## Introduction

Intravenous Immunoglobulin’s (IVG’s) are prepared from up to 60,000-pooled plasmas and highly efficacious therapeutics used for an increasing variety of medical conditions (Harvey 2005, Miescher et al., 2005). Clinically IVIGs are safe, generally well-tolerated and highly efficacious treatments for an ever-expanding array of medical conditions that include, primary and secondary immune deficiencies, autoimmunity, inflammation, and peripheral neuropathies (Stiehm 1996, Simon and Spath, 2003, Looney and Huggins 2003). Experimental and clinical evidence reveal that IVIG work in a variety of ways, and these often multifactorial, but the exact mode(s) of action for many conditions unknown (Harvey 2005). Traditionally Australia had been self-sufficient for its internal sourcing and supply of blood-derived products, including IVIGs. However the AUSFTA (the Australian – United State Free trade Agreement) operational from January the 1^st^ 2005 opened the door for the importation and use of foreign (overseas) blood products for Australian clinical use (AUSTFA 2006, Bambrick et al., 2006). Thus prompting fundamental questions as to whether locally sourced and prepared IVGs in Australia were both physically and biologically identical with their counterpart products extracted and purified outside of Australia. This present study addresses a proteomic comparison of IVIGs whist a separate and parallel study was performed to compare the biological similarities and differences between IVIGs extracted from different global populations for antibodies to human and avian influenzas (Lynch et al., 2006a). For both studies the central question being asked was whether the various formulations were effectively equivalent, or actually notably and significantly different?

Effectively IVIGs are mega-clonal mixes of broadly reactive Antibodies (Abs) that reflect the broad herd immunity of the respective source population(s) and the antigenic environments of those locales. Comparative fine-detail proteome mapping and comparison of IVIG formulations prepared from Australian versus overseas plasma sources had not been presented. This was considered particularly important since the various foreign-derived immunoglobulin formulations now being used in Australia (a highly and uniquely antigenic island continent), are from distinctly different genetically and geographically blood-donor populations, and those antigenic (Ag) environments, than the Australian populace, and its flora and fauna. Which, including product preparation differences led to the prediction that Australian derived and non-Australian derived IVIG formulations may differ in antibody compositions, functional target specificities or additional carry-over proteins. Added to this the IVIG formulations studied were extracted and purified by different methods (Nydegger and Mohacsi 2002, Martin 2006) and yield different levels of product purities of 94 to 98 % and greater. With this in mind it was considered highly likely the different IVIG formulations would differ from one another in physical and compositional properties (this study) and possibly also in biological activities (Lynch et al., 2006a). With the belief that detailed and comparative understanding of the various IVIG formulations, their similarities and differences in biochemical characteristics and bio-specificities, was needed, as was knowledge of whether any other non-immunoglobulin proteins might have been carried along from plasma through processing and purification. The underlying premise being that with enhanced understanding of the proteomes of IVIG therapies might enable a more strategic and optimal use of formulations, and possibly some identified to be ideally tailored for strategic therapeutic use for specific conditions.

Further to this, to better understand the molecular basis of some of the common well noted but extensive list of side effects of IVIG infusion, including headaches, nausea etc., that can be experienced (Pierce and Jain 2003, Orbach 2005, Gelfand 2006), in this study we investigated evidence of possible carry-over cytokines across a number of IVIG preparations sourced from different geographical locales. Hence, driven by a need for better understanding of IVIGs, their compositions and antibody specificities, and questions of similarities and differences, led to the hypotheses; that i) IVIGs prepared from geographically different donor population may notably differ in their antibody characteristics and non-immunoglobulin components, and ii) that Proteomic profiling of IVIG formulations may aid in a better molecular understanding of IVIG mechanisms of action and treatment adverse reactions.

In this study of IVIGs, a valuable therapeutic resource that has increasingly been difficult for supply to keep up with its demand we have used proteomics to broadly compare the respective protein compositions of IVIGs from different geographical populations. There is no question that biologically IVIGs from one country to another are highly effective and valuable therapeutics wherever they are used. But so saying, IVIG cannot be directly considered the same as for example as a chemical drug, which if prepared in exactly the same manner, using exactly the same staring chemicals and solutions will yield exactly the same products of absolutely identical molecular structure and function. IVIGs in comparison are derived from thousands of blood plasmas, which even from a single person depends on the genetics, geography, climate, environmental antigens, diet, physical health, health complication, sex, age, season of the year, prior infectious agent exposures, travel history etc., of that individual. So for a single product those differences are multiplied even further by each of the separate individual donations used for a single preparation (i.e., batch). And further again, across multiple time-points and global population sampling. Adding yet further to these inherent natural differences, a variety of different methods are used in preparation of IVIGs. As a result the immunoglobulin products themselves vary in their levels of immunoglobulin purity, equal to or greater than 95%.

Hence our starting hypothesis was that the various IVIG formulations might vary notably in composition and perhaps activity. Therefore to test the first of these our study aimed to; *a*) assess and compare the individual proteomes of the respective IVIGs derived and prepared from the different geographical donor populations, and *b*) Identify and characterise any specific differences between preparations, and *c*) using proteomics identify any non-immunoglobulin proteins carried along from plasma through the immunoglobulin purification process, and *d*) in more targeted approach, use multiplex xMAP analysis of the IVIG preparations, to detect the presence of any co-isolating cytokines.

In brief for this study we have used a proteomic approach to characterise and compare the various IVIG formulations clinically used in Australia, and so gain a more definitive understanding of their compositions and equivalence. The findings of these studies did reveal notable differences between IVIG preparations, in the global characteristics of the Immunoglobulin components, variability in carry-over other plasma proteins, and in particular, the presence of cytokines of different types and levels, in and between formulations. Whether such differences are of biological or functional consequence remains an intriguing but unknown and unresolved point of speculation.

## Materials and Methods

### Intravenous Immunoglobulin

For these studies highly purified IVIG formulations were extracted from American donors (Gammunex, Grifols, Clayton, North Carolina/Los Angeles, California)(Carimune, ZLB Biologicals, Berne, Switzerland) (Octagam-US, Octapharma, Vienna Austria), US-European donors (Octagam, Octapharma, Vienna, Austria), European donors (Sandoglobulin, CSL-ZLB, Bern Switzerland), Australian donors (Intragam P, CSL Ltd, Melbourne Australia), and South East Asian donors (Intragam P-SE, CSL Ltd, Melbourne Australia) pooled blood donations. Manufacturing differences of the respective products was reflected in the listed purities of those products, with the product sheets for the Intragram preparations listed at purities of > 98%, and the others at ≥ 95%. Because many of the formulations differed in starting product concentration these were diluted in sterile saline as required for protein equivalence in testing.

### Electrophoresis and Mass Spectrometric Profiling of IVIGs

*Electrophoresis*: The 2D Electrophoresis and Mass Spectrometry studies were performed in collaboration with Minomic Pty Ltd. (www.minomic.com) (Lynch et al., 1996, Nouwens 2000, Lynch et al., 2006b). Basically 300 ug of IVIG protein were subjected to isoelectric focusing (IPG) across pH gradients of 4-7 (Biorad, LifeSciences, Hercules, Ca, USA) or pH 6-11 (GE Healthcare, Uppsala, Sweden). With IPG strip reduction, alkylation, and detergent exchange, and 2D-sodium dodecyl sulfate-polyacrylamide gel electrophoresis (SDS-PAGE) replicates performed on 11cm 8-16% (w/v) polyacrylamide/Tris-HCl gels (Bio-Rad) using a Criterion Dodeca Cell (Bio-Rad). Gels were fixed in 10% (v/v) methanol, 7% (v/v) acetic acid, and stained with Sypro Ruby (Molecular Probes, Or, USA), and over-stained with Colloidal Coomassie blue G-250). The gels were de-stained and scanned using a ProXPRESS Proteomic Imaging System (PerkinElmer, Ma, USA).

For analysis of low molecular weight protein fragments in IVIG preparations including evidence for any non-immunoglobulin protein carried along through the IVIG separations from plasma, size exclusion fractionation was performed. For this molecular weight cut-off filters (Millipore Filters (Sigma-Alrich, St Louis, Mo, USA)) were used to capture proteins and protein fragments in the less than 24 kDa flow through fraction. These were concentrated, trypsin digested and peptides resolved by LC-MS at Minomic Pty Ltd www.minomic.com). From database searches the respective identities of the recovered peptides were identified and these listed in table-1.

### Mass Spectrometry

Mass Spectrometry was performed on the respective IVIG preparations to firstly broadly assess the biochemical properties of the immunoglobulins within them, as well as any other plasma proteins or protein fragments that may have been carried along through the respective purification procedure. For this gel plugs were cut from the 2-D gels, destained and protein digested overnight at 37°C with trypsin (Promega, Madison, Wisconsin). The digested peptide samples were desalted and concentrated using C18 zip tips (Eppendorf, GmbH, Hamburg, FRG) and spotted onto MALDI-MS target plates with a-cyano-4-hydroxycinnamic acid. Samples were then analysed by MALDI-TOF MS (Waters, Milford, Ma, USA), run in reflectron mode. The recovered data was searched over the Swissprot database (www.uniprot.org) using the Mascot search engine (www.matrixscience.com).

### Multiplex Analysis of Cytokines in IVIG preparations

Analysis for cytokines in IVIG’s was performed by Multi-Analyte-Profiling (xMAP) assays out-sourced to Upstate Biotechnology (Lake Placid, NY, USA) and Rules based Medicine (Myriad-RBM, Austin, Texas, USA), using 22/26-plex Human cytokine arrays, with each cytokine demarcated to micro-beads of a uniquely coloured fluorophore for identification and standard curves. Tests, controls and standards reactions were read using a Luminex_100 biosystem (Luminex, Austin, Texas, USA). These xMAP assays were performed as described previously (Rhyne et al., 2003).

For the Beadlyte^®^ multiplex assays (Rhyne et al., 2003) performed at Upstate (www.upstate.com) for the detection of cytokines in IVIGs formulations prepared from European, South-East Asian or European-US blood donations are presented in Figure 2 as H, I, J. Cytokine levels are compared against those of 2 normal samples of Human serum. Separate evaluations of other panels of an Australian and 3 United States (US) derived IVIG preparations (listed as formulations K, L, M, N) was also performed at Rules Based Medicine (www.rulesbasedmedicine.com). In this instance cytokine values are compared against established high-range human serum levels for that cytokine.

**Figure 1:**
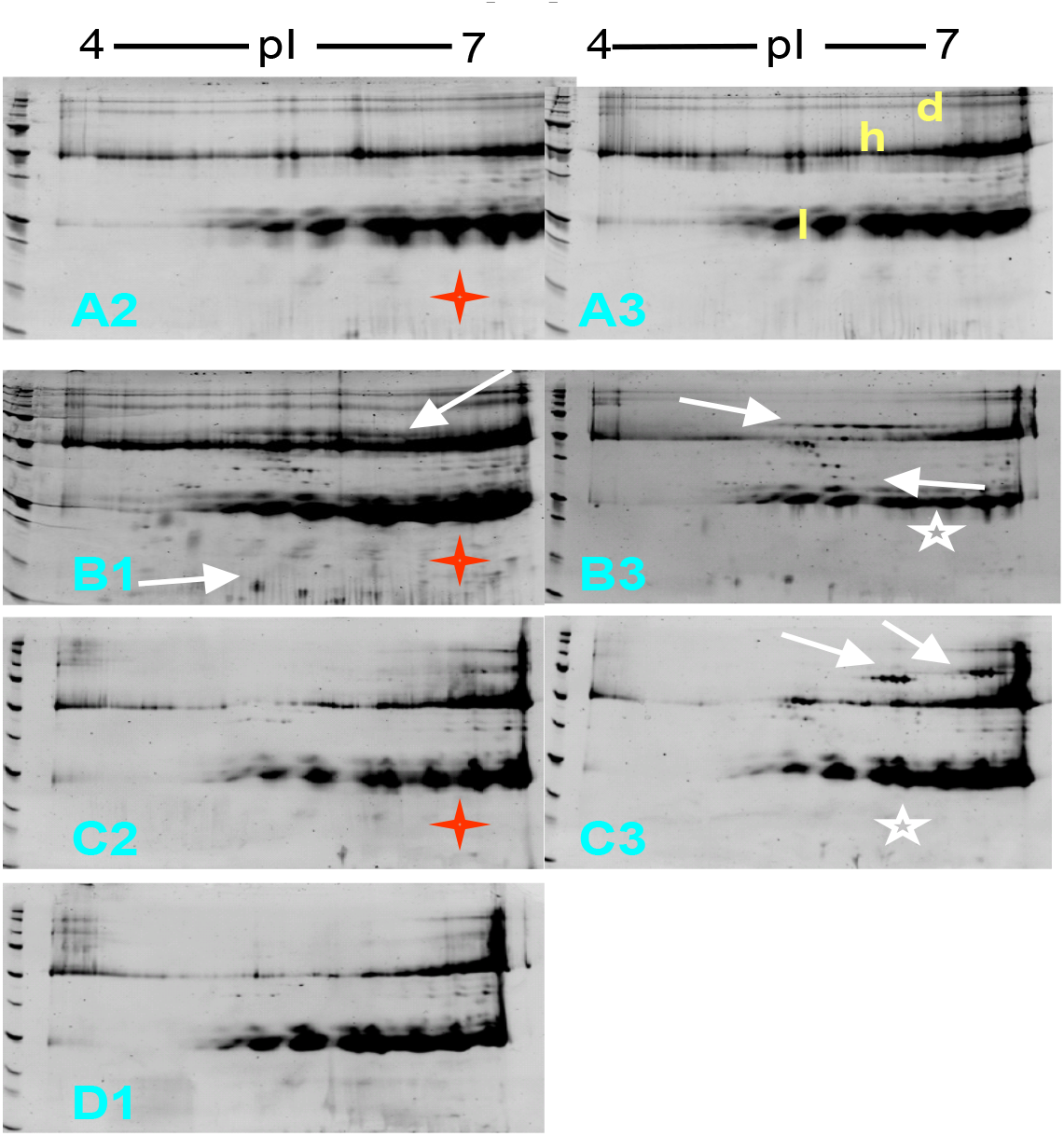
2D-electrophoretic profiling of 4 different (A-D) IVIG formulations of Igs prepared from Australian, European and North American donated plasmas. IVIGs (with different batches for A-C) were subjected to isoelectric IPG separations using 4 to 7 (also 6 to 10, data not shown), electrophoresis and protein stain (Sypro Ruby). Indicated are the light (l) and heavy (h) chains of the immunoglobulins and Ig dimers (d).

**Fig 2.**
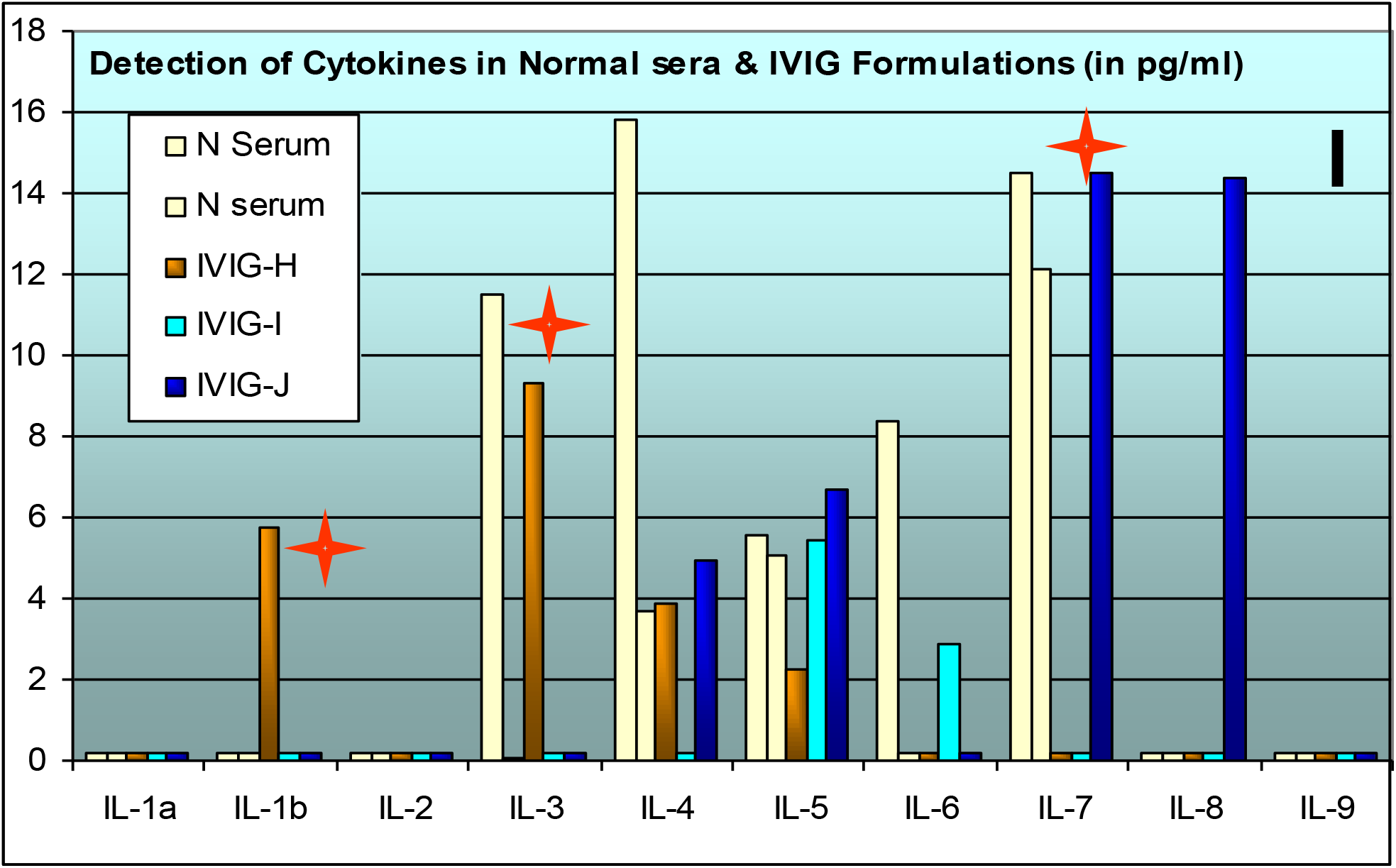
Use of Upstate-(xMAP) Multiplex Assay for Cytokines in IVIG Preparations. Shown at the top (panel I) are the cytokines detected by multiplex assay as compared with 2 normal sera in representing European/ Sth-East Asian and European-US donor formulations (ie randomly assigned and plotted as H, I, J) and assayed commercially by Upstate (www.upstate.com) using Beadlyte multiplexing.

**Fig 3.**
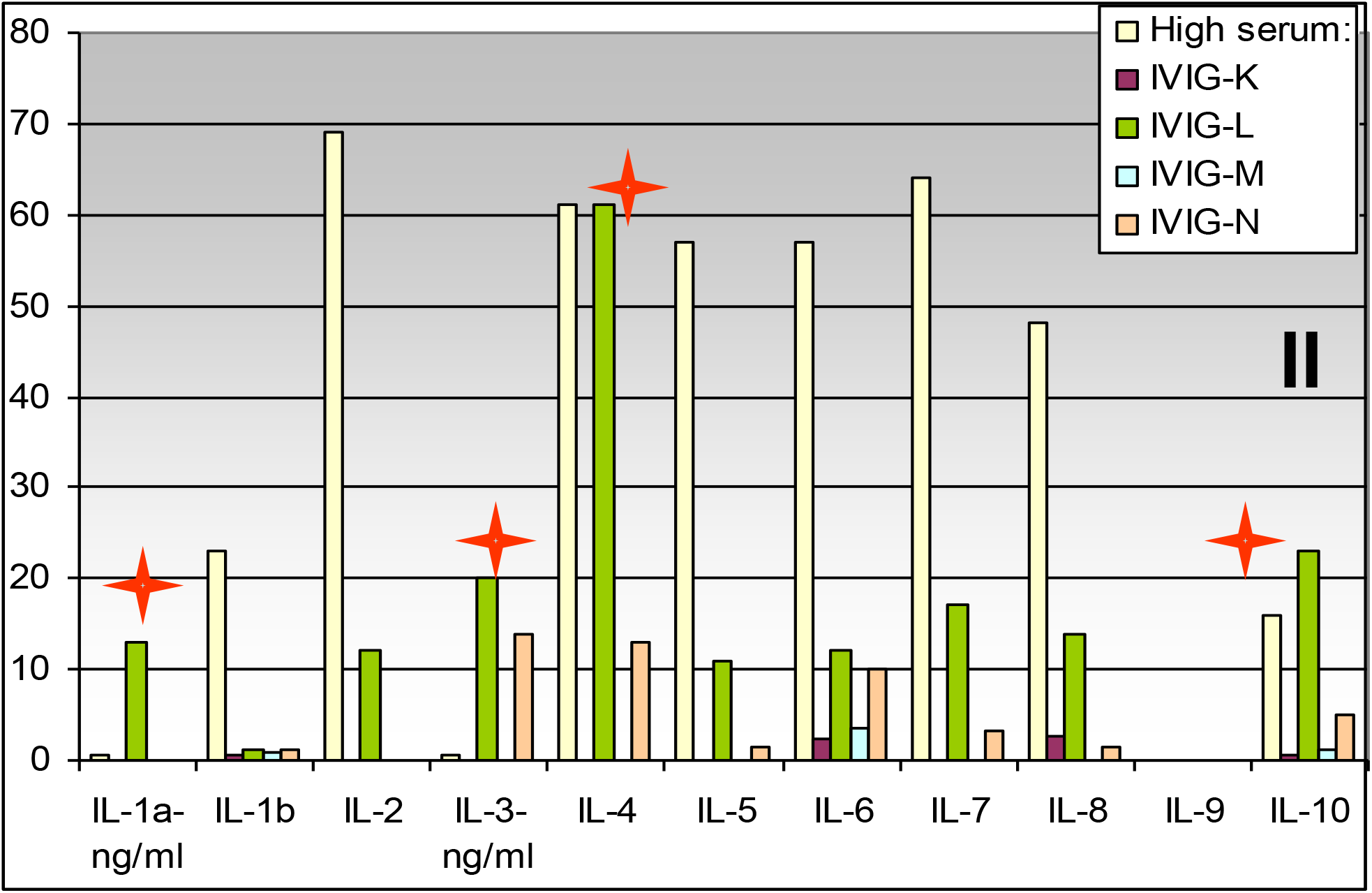
Myriad RBM -(xMAP) Multiplex Cytokine Assay of IVIG Preparations. An Australian-derived IVIG preparation and 3 US-derived IVIG formulations (ie K-N) were commercially assayed by Rules based Medicine (www.rulesbasedmedicine.com) and the cytokine values obtained, plotted against the high range of levels found in human serum. The red stars identify levels of the respective cytokine that are equivalent to or greater than levels found in plasma.

## Results and Discussion

### i) 2D-Gel Electrophoretic Profiling of IVIGs

Despite similar global pI spreads in the Ig components of the different preparations there are notable differences between IVIG preparations in the monomer and dimer fraction and in particular with non-Ig proteins additional to the Ig protein trails (Figure 1)(some of those differences are highlighted by arrows). Overall the higher purity of the Australian preparation (A) as listed at >98% Ig compared to the overseas preparations (B, C) listed at > 95%, is shown here, and with a greater non-Ig protein component (indicated by arrows, and white stars) identified in the latter. Variance in staining profiles between batches of the same product are also indicated, e.g., with the respective batches labelled 1-3.

The profiles of the low-molecular weight proteins in preparations A, B and C revealed antibody fragmentation and a number of non-immunoglobulin proteins. These extra proteins were more extensive in the overseas formulations than in the Australian preparation as also identified in this fraction (indicated by arrows) of the Figure 1, protein stained maps.

As proteomic maps of IVIG preparations were not available in the public domain the Proteomes of the IVIGs used in Australia were mapped and it became evident that, there were considerable differences between the proteomes of the different formulations. These findings add to a growing list of other studies that highlight that IVIGs differ in a range of clinical and other characteristics (Gelfand *et al*., 2006) and reinforced the proposal that IVIGs isolated from different (geographically separate) donor populations and prepared using a variety of different procedures, were not identical. In comparing formulations marked qualitative differences in the respective 2D-gel Ab profiles were identified. These include differences in Ab pI spreads, monomer/dimer ratios and in some cases multiple, molecular weight (MW) Ig species. The underlying molecular basis for the latter is of particular interest as it is likely that differences in purification methodologies may select or deselect for Abs with differing properties. Hence ongoing studies to assess the post-translational glycosylation and thiol properties of the respective preparations are warranted.

Mass spectrometric identification of non-Ig components carried along from plasma through purification was performed on low MW proteins fractionated by size-exclusion filtration. Few ancillary proteins or Ab fragments have so far been identified in Australian Sourced and prepared IVIGs (listed at >98% purity). This contrasted to the overseas preparations (≥ to 95% purity) that revealed several carry-over proteins, including; complement Factor-B, serotransferrin, and VEGFR1 proteins, as well as heavy and light chain Ab fragments.

### ii) Detection of Cytokines (IL1 – IL10) in IVIG Formulations by Muliplex (MAP) Assay

In an alternate strategy IVIGs were subjected to commercial multiplex analysis for direct evidence of cytokines carried over from the pooled plasmas in any of the IVIG formulations under study. For this, IVIGs were assayed at Myriad-RBM or Upstate Biotech which both use highly validated, Luminex-based multiplex xMAP (Multi-Analyte-Profiling) bead assays, for the detection of Human cytokines (Carson and Vignali, 1999, Elshal and McCoy, 2006). Cytokines are small molecular weight (MW) proteins of 3-24 kilodaltons, and highly active at low pico to nano gram per ml quantities (Dinarello 1999), and central to diverse tissue functions and both positive and negative mediators for many clinical conditions and disease states.

As noted there are considerable differences in the cytokines measured between the different preparations as there are in the levels expressed in single normal sera. These differences are representative of a wider examination of cytokines, growth factors and chemokines (data not shown) between preparations commercially tested by fluorescence multi-analyte bead Rules Based Medicine and Upstate Bioscience Assays. As expected findings to-date within product variations are less, however, inter batch variability is apparent and consistent with differences between levels in normal sera.

#### Upstate xMAP Studies

#### Myriad-Rules Based Medicine xMAP Studies

The importance of these findings of evidence of different cytokines in some IVIG preparations is that it further supports the conclusion of notable variance in non-immunoglobulin protein composition between the different IVIG formulations. With this said it is essential to note that it does not clarify in any way on the source of this material. That is whether it was co-purified as unbound cytokine carried through the extraction procedures and, or whether it actually represents bound-cytokine captured by naturally derived autoantibodies in the respective IVIG preparations. So whilst these results clearly indicate the variable presence of cytokines in some IVIG products, at this point they do not elaborate as to whether the detected cytokine(s) represent free-unbound cytokine protein, or antibody-bound cytokine that may or may not be naturally dislodged, or released by the treatment conditions. But most importantly these findings in themselves provide no information as to whether such detected cytokine material is bioactive. Hence it must be stressed that these findings merely open a window of consideration that *<u>if</u>* such cytokine in IVIGs were biologically active they *<u>may</u>* potentially have some positive or negative biological effects. And if so the latter would predictably depend on the clinical condition for which the IVIG was being used. Up-to this point, this is merely conjecture, but none-the-less an intriguing possibility.

Collectively, these findings reveal significant differences between the compositions and biochemical characteristics of locally derived and overseas IVIG preparations. This suggests caution in attributing equivalence between IVIG formulations, at least until there is a detailed mining and comparative understanding of their biochemical properties and Ab specificities, to be matched with clinical use. Detailed IVIG characterizations thus may; i) enable the development of Ab enriched/ fractionation strategies to select functionally specific Abs of set biochemical characteristics, and thus ii) facilitate specific and streamlined use of particular IVIG preparations for different and select clinical targets and conditions (e.g., autoimmune disease vs infectious agent Ags).

## Conclusions

The proteomic maps of several batches of each of the different IVIG formulations used in Australia have been examined. As expected this revealed considerable similarity between products. But it also elaborated a variety of quite notable differences, which included remarkable variance in the monomer to dimer Immunoglobulin fractions and, or presence of non-immunoglobulin proteins across the formulations. Using mass spectrometry it was possible to directly identify evidence for a number of non-immunoglobulin carry-over proteins (and/or fragments) in the IVIGs (i.e., Table 1: Caspase 3 precursor, VDJ recombination activating protein 1, RAG 1, Gelsolin, Apolipoprotein 1, NADH ubiquinone oxidoreductase, Complement factor B precursor, Serotransferrin, VEGF receptor 1 precursor, Tetranectin precursor). As might be predicted the preponderance of the non-Ig proteins were in formulations with listed purification levels at ≥ 95%.

**Table 1:**
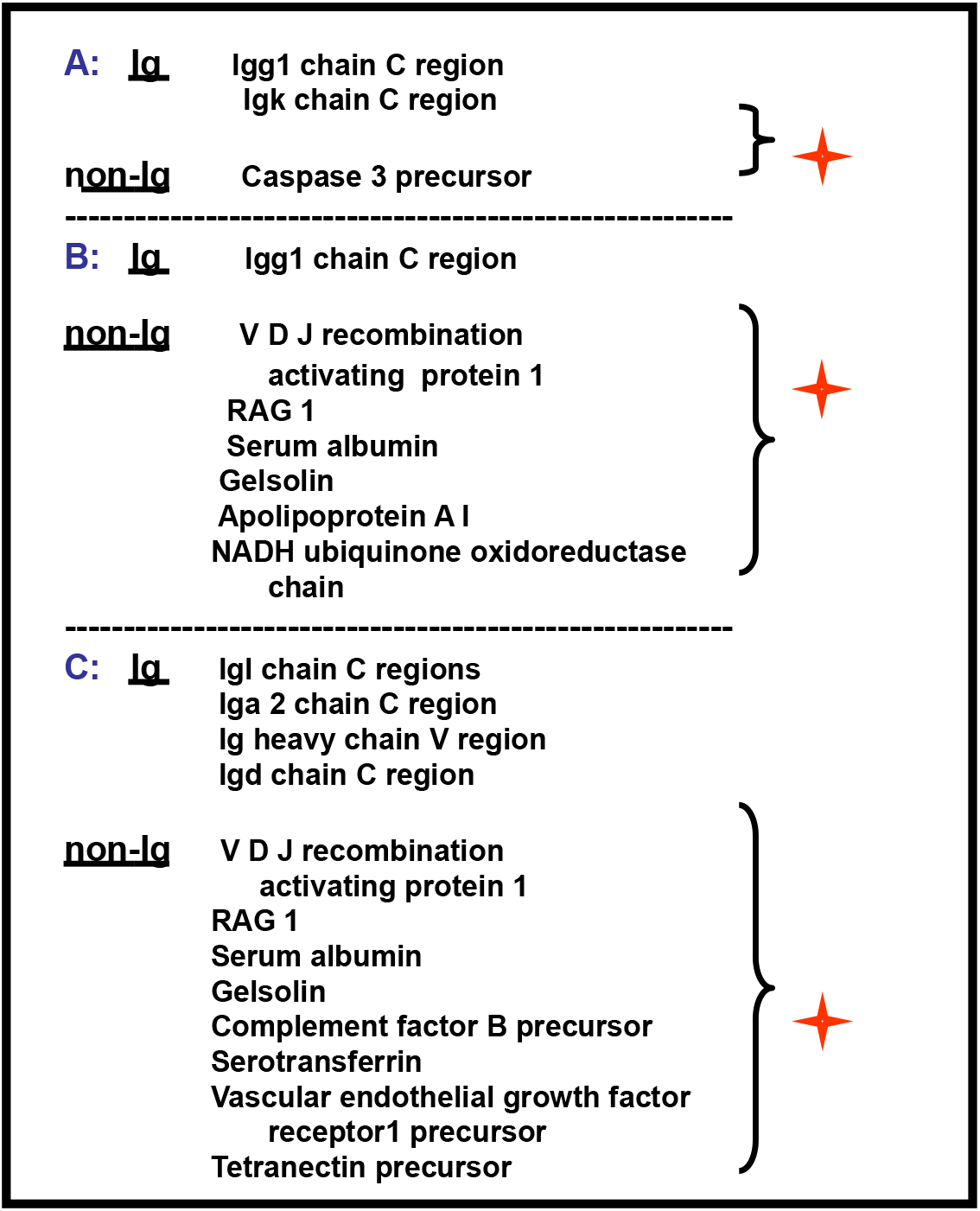
Mass Spectrometric Identification of proteins fractionated from the different preparations by 24 kDa, filter exclusion. The flow through fraction of IVIG preparations fractionated using 24 kDa cut off filters (i.e., proteins <24 kDa), were concentrated, trypsin digested and peptides resolved by LC-MS (at Minomic Pty Ltd). The protein identities were made from Mascot database searches.

In a completely different approach additional variability in non-immunoglobulin protein in and across the IVIGs was observed, with multiplex-assay evidence of cytokines in each of the IVIG formulations using 2 different commercial multiplex studies (Figure 2). The unequivocal conclusion from this is that there is evidence for the presence of some cytokines in IVIG preparations, and that their detection differs across products. The important next step is to determine whether these differences are of therapeutic relevance, either as positive or negative effectors of IVIG infusion or IVIGs delivery of clinical benefit for the specific medical conditions they are being used to treat. Of the batches studied to date the Australian produced batches were found to be the most homogeneous and as listed were of highest Immunoglobulin purity. In contrast the proteomes of the overseas formulations studied were found to contain higher non-immunoglobulin protein and greater differences in their Immunoglobulin profiles between batches. It will be important to determine whether detection of cytokines in IVIGs can be profiled and correlated with the nausea, headaches etc., side-effects observed following some treatments, and whether specific cytokines in some IVIG preparations may positively or negatively act in provision of clinical benefit.

Core to the demonstration in these studies that have revealed both similarities and differences in the physical and potential biological activities within and between batches of IVIGs was a far greater realisation of each formulations potential to be a reflective map of the overall herd immunity of the donor source population. Given that such immune responses would themselves be routed in the respective environmental antigens and circulating pathogens of the source population and time of collection, as well as the overall health of that population. And a correlate of this, was a growing belief that IVIGs could offer a tangible means as a biological sentinel to one assess and map pathogens, and a populations response to them. And one that would effectively include a temporal historical and seasonal mapping of both strong dominant, as well as broad subdominant, antibody immune responses of a community exposure to persistent as well a spasmodic environmental pathogens. As most efforts on pathogen targeting have historically been from a specific antigen targeting for discrete highly pathogenic strains of a select infectious agent, asked was whether this unique situation of IVIG could be explored to ask whether humans have broad immune (antibody) capabilities that are directed across all strains, not just select ones of a pathogen. And if so whether such activities could offer a degree of protective control, against both mild as well as highly virulent forms of a pathogen. From such realisations in 2004 it was predicted that indeed such broadly protective antibodies do exist, and that IVIGs held the unique key to proof of this existence, as well as the molecular specificities of such antibodies, and therefore highly unique capacity for exploration of cross-reactive, pre-existing, broadly cross-reactive antibodies for infectious diseases. Our studies flowing from this over the intervening almost two decades were initially focussed on Influenza as the pathogen of study, and later expanded to a wider variety of other viruses and a bacterium. With the basic aim that by understanding and mapping the broad strain immune targets of pathogens by IVIG antibody profiling would enable firstly the realisation of antibodies in an existing accepted and well tolerated therapeutic, for potential use in passive immune protection. And additionally provide a key to revealing the molecular signatures on pathogens for strain specific as well as cross-strain acting antibodies, for more universal and broadly acting vaccines and other therapies.

It is hopeful that the findings of studies performed and presented on Immunoglobulin formulations of nearly two decades ago, might now prompt new considerations of current IVIG preparations. And perhaps guide thoughtful new study approaches of the proteomes of present-day IVIGs, and to possibly include activity-based analyses to value-add to IVIG transfusion treatments and management.

## Acknowledgements

The title page lists the current addresses of investigators. The studies described were performed across the Australian Red Cross Blood Service, Sydney University and Minomic over the 2004 to 2006 period, and supported for IVIG Proteome studies by the ARCBS and the University of Sydney. NK was the recipient of a Sydney University Summer Student Research Scholarship. We thank Dr Y Ayob for study material access. Presented are findings on the composition and activities of IVIGs as presented at the 3^rd^ Australian National Health and Medical Congress (www.ahmrcongress.au), held at the Melbourne Convention Centre, 26^th^ November - 1^st^ of December 2006, Melbourne, Victoria, Australia. A copy of the poster presentation from that conference is attached as Supplement Figure S1, and includes the respective investigator address details of that period. Far more complete, detailed and referenced accounts of this work were also presented as project grant applications including through the 2005-2006 period, requesting Government grant support via the National Health and Medical Research Council (NHMRC) of Australia, submission system. One such project application (i.e., NHMRC_ID-402877) was titled: ‘*Determine the Anti-Influenza Ab Component of IVIGs (Intravenous Immunoglobulins) Clinically Used in Australia: Proteomic & Virologic Assessment of anti-influenza IVIg potential’* and originally submitted as an EOI in November 2005, in response to a NHMRC Urgent Research Grant call ‘for Research into a Potential Avian-Influenza-Induced Pandemic’, and, follow-up request for full application; G Lynch (CI), J Sullivan, et al., in March 2006. Further to this, the findings herein the article and in related studies were used to advise and inform on IVIGs for a special subcommittee (that included Sir Peter Lawler (OBE) and Professor Kevin Rickard (AM)) of the 2006 Review of Australia’s Plasma Fractionation Arrangements (Flood *et al*., 2006). The observations made were also the starting point and backdrop of IVIG studies and 11 NHMRC grant submissions over the ensuing decade. With studies that collectively underpinned the existence, realisation, and definition of a unique and separate under-appreciated area of adaptive immunity, and for which its guiding molecular interactions have been defined in a separate ‘theory of dark immunity’^©^ by authors GL and JS.

## Disclaimer

As acknowledged in the text these historical proteomic and multiplex studies for IVIGs prepared around the 2001 – 2006 period, were compositional and not function or activity-based. Furthermore because of processing advances and method differences over the interim period the findings made cannot therefore be directly equated to current 2021 era IVIG formulations, which would need to be freshly assessed on their own merit.

