## Supplemental Figure S1 for "Translational Proteomics for Transfusion Medicine: Resolution of the IVIG Proteomes of Different Geographically Sourced and Prepared IVIG Immunotherapies"

**Supplement Contents Overview;**

Supplement page Numbers:

**SM1**: Article Title Information and Supplement Contents Overview.

**SM2**: Supplement Figure S1 of Poster presented in 2006 of the article studies, namely; Lynch, G.W, Fitzgerald, A., Walsh, B., Kapitza, N., J. S. Sullivan, J.S. (2006) Translational Proteomics for Transfusion Medicine: Resolution of The IVIG Proteomes of Different Geographically Sourced and Prepared IVIG Immunotherapies. 26th Nov – 1st Dec. 2006. Melbourne, Victoria. *Australian Health and Medical Research Congress Proceedings.* Volume **5**, 2006

**SM3**: *2005-2006 Conference listings of these and closely related studies:* &
*Historic background of our Proteomic and Influenza-based studies of IVIGs, and wider implications:*

**SM4**: *Supplement References:*

### Translational Proteomics for Transfusion Medicine: Resolution of the IVIG Proteomes of Different Geographically Sourced and Prepared IVIG Immunotherapies.

Australian Red Cross
BLOOD SERVICE

**G. W. Lynch[1,2], A. Fitzgerald[3], B. Walsh[3], N. Kapitza[1,2], J. S. Sullivan[1,2]**

*[1]Research and Development, Australian Red Cross Blood Service, Sydney, NSW, [2]Transfusion Medicine & Immunogenetics, Faculty of Medicine, University of Sydney, Sydney,NSW, [3]Minomic Pty Ltd, Chatswood West NSW, Australia*

**

minomic

**Abstract:** Intravenous Immunoglobulins (IVIG) are prepared from up to 60,000 pooled plasmas and comprise a megaclonal mix of Antibodies (Abs) of broad specificity. Clinically IVIGs provide generally well tolerated and highly efficacious treatments for primary and secondary immune deficiencies, auto-immune conditions, inflammation, and peripheral neuropathies. Experimental and clinical evidence indicates that IVIG work in a variety of ways, but the exact mechanisms of action for most conditions are unknown. Furthermore, comparative fine-detail proteome mapping of IVIG formulations used in Australia, but isolated from various local and overseas plasma sources, have not been performed. As the different formulations are from genetically and geographically distinct populations and antigenic (Ag) environments, and prepared using different isolation methods, they predicably would differ. We hypothesise that a detailed comparative understanding of each of the formulations, their biochemical characteristics and bio-specificities, will permit an optimal and efficacious matching for specific target Ags and conditions. When comparing formulations we have identified marked qualitative differences in their respective 2D-gel Ab profiles. These include differences in Ab pI spreads, monomer/ dimer ratios and in some cases multiple molecular weight (MW) Ig species. The underlaying molecular basis for the latter is of particular interest as it is likely that differences in purification methodologies may select or deselect for Abs with differing properties. Hence ongoing studies to assess the post-translational glycosylation and thiol properties of the respective preparations are warranted. Mass spectrometric identification of non-Ig components carried along from plasma through purification was performed on low MW proteins fractionated by size-exclusion filtration. Few ancillary proteins or Ab fragments have so far been identified in Australian Sourced and prepared IVIGs (listed at >98% purity). This contrasted to the overseas preparations (≥ to 95% purity) that revealed several carry-over proteins, including; complement Factor-B, serotransferrin, and VEGFR1 proteins, as well as heavy and light chain Ab fragments. In an alternate strategy IVIGs were subjected to multiplex analysis revealing a variety of different cytokine profiles between the formulations. Collectively, these findings reveal significant differences between the compositions and biochemical characteristics of local and overseas IVIGs. This suggests caution in attributing equivalence between IVIG formulations, until there is a detailed mining and comparative understanding of their biochemical properties and Ab specificities matched with clinical use. Detailed IVIG characterizations thus may; i) enable the development of Ab enriched/ fractionation strategies to select functionally specific Abs of set biochemical characteristics, and thus ii) facilitate specific and streamlined use of particular IVIG preparations for different and select clinical targets and conditions (eg autoimmune disease vs infectious agent Ags).

#### Hypotheses

- **IVIGs prepared from geographically different donor population differ significantly in their antibody characteristics and non-immunoglobulin components**
  - **Proteome mapping of IVIG formulations will aid our molecular understanding of treatment side-effects, adverse reactions and mechanisms of action**

#### Aims

- **Globally assess the IVIG proteome of preparations derived from different geographical donor populations**
  - **Identify and characterise specific differences in the proteins of different preparations**
    - **Identify the non-immunoglobulin components of IVIG preparations**
      - **and in particular determine whether IVIGs contain cytokines**

**Introduction:** IVGs are highly efficacious therapeutics for a wide and increasing variety of medical conditions and worldwide their provision is a billion dollar industry. Surprisingly the mechanisms of action of IVIGs are increasingly being appreciated as multifactorial, and for most conditions have not been resolved. Furthermore the molecular basis of common side effects (up to 30% in some populations) have not been identified. We ask whether there are carry-over proteins in IVIGs that are isolated along with the antibodies, as these may contribute to infusion side effects (headaches , nausea etc). IVIGs for their protein compositions and antibody specificities is necessary to better understand their similarities and differences and their use for a wide variety of medical conditions. Better understanding of the proteomes of IVIG is required to enable optimal targeting and use of the respective formulations for specified conditions. IVIGs are a valuable therapeutic resource but one which is increasingly difficult for supply to keep up with demand. In this respect we ask what are the different protein compositions of IVIG preparations clinically used in Australia.

#### 2D-Gel Electrophoretic Profiling of IVIGs

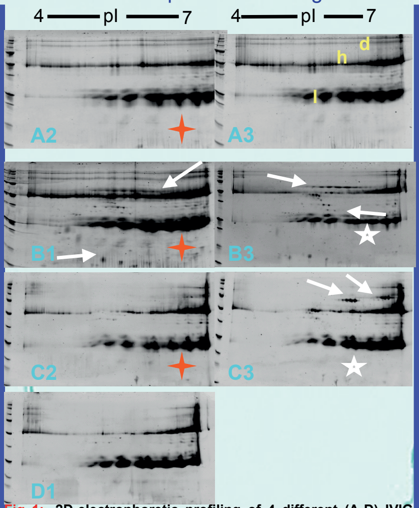


**Fig 1: 2D-electrophoretic profiling of 4 different (A-D) IVIG formulations of Igs prepared from Australian, European and North American donated plasmas.** IVIGs (with different batches for A-C) were subjected to isoelectric IPG separations using 4 to 7 (also 6 to 10, data not shown), electrophoresis and protein stain (Sypro Ruby). Indicated are the light (l) and heavy (h) chains of the immunoglobulins and Ig dimers (d).

**Results**: Despite similar global pI spreads in the Ig components of the different preparations there are notable differences between IVIG preparations in the monomer and dimer fraction and notable differences in the apparent non-Ig proteins additional to the Ig protein trails. These differences are highlighted by arrows. Overall the higher purity of the Australian preparation (A) listed at >98% Ig compared to the overseas preparations (B,C) listed at > 95%, is shown here with greater non-Ig protein and differences (indicated by arrows, and white stars) identified in the latter. Differences between batches are also highlighted

**A:** **Ig**
- **Igg1 chain C region**
- **Igk chain C region**

**non-Ig**
- **Caspase 3 precursor**

---

**B:** **Ig**
- **Igg1 chain C region**

**non-Ig**
- **V D J recombination activating protein 1**
- **RAG 1**
- **Serum albumin**
- **Gelsolin**
- **Apolipoprotein A I**
- **NADH ubiquinone oxidoreductase chain**

---

**C:** **Ig**
- **Igl chain C regions**
- **Iga 2 chain C region**
- **Ig heavy chain V region**
- **Igd chain C region**

**non-Ig**
- **V D J recombination activating protein 1**
- **RAG 1**
- **Serum albumin**
- **Gelsolin**
- **Complement factor B precursor**
- **Serotransferrin**
- **Vascular endothelial growth factor receptor1 precursor**
- **Tetranectin precursor**

**Table 1: Mass Spectrometric Identification of proteins fractionated from the different preparations by 24 kDa filter exclusion.** The flow through fraction of IVIG preparations fractionated using 24 kDa cut off filters (ie proteins <24 kDa), were concentrated, trypsin digested and peptides resolved by LC-MS (at Minomic Pty Ltd). From data-base searches the identities of the recovered peptides were revealed and listed in table-1. **Results**: The profiles of the low-molecular weight proteins in preparations A, B & C revealed antibody fragmentation and a number of non-immunoglobulin proteins. These extra proteins were more extensive in the overseas formulations than in the Australian preparation as also identified in this fraction (indicated by the red arrows) of the Fig 1 protein stain maps.

#### IVIG Ab-Muliplex Ag-Binding Comparison

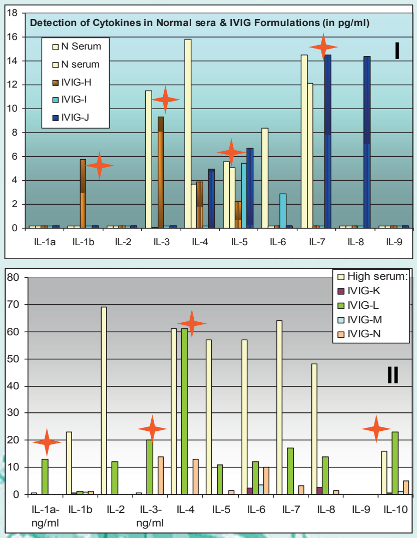


**Fig 2. Use of Multiplex Assay for detection of Cytokines in IVIG Preparations**. Shown in top panel I are the cytokines detected by multiplex assay as compared with 2 normal sera in representing European/ Sth-East Asian and European-US donor formulations (ie randomly assigned and plotted as H, I, J) and assayed commercially by Upstate (www.upstate.com) using Beadlyte multiplexing. Similarly an Australian and 3 US preparations (ie K-N) were commercially assayed by Rules based Medicine (www.rulesbasedmedicine.com) and the values obtained plotted against the high range of levels found in human serum. The red stars identify levels of the respective cytokine that are equivalent to or greater than levels found in plasma. **Results:** As noted there are considerable differences in the cytokines measured between the different preparations as there are in the levels expressed in single normal sera. These differences are representative of a wider examination of cytokines, growth factors and chemokines (data not shown) between preparations commercially tested by fluorescence multi-analyte bead Rules Based Medicine and Upstate Bioscience Assays. As expected findings to-date within product variations are less, however, inter batch variability is apparent and consistent with differences between levels in normal sera.

**Discussion:** As proteomic maps of IVIG preparations are not available in the public domain we have mapped the Proteomes of the IVIGs used in Australia and show that there are considerable differences between the proteomes of different formulations. These findings add to a growing list of other studies that highlight that IVIGs differ in a range of clinical and other characteristics (Gelfand et al 2006) and reinforces that IVIGs isolated from different donor populations, often geographically different and prepared using different procedures, are not identical. Of the batches studied to date the Australian batches were found to be homogeneous, in contrast the proteomes of the overseas formulations studied were found to contain higher non-immunoglobulin protein and greater differences in their protein profiles between batches. It will be important to determine whether specific cytokine profiles correlate with the nausea, headaches etc side-effects observed following some treatments, and whether specific cytokines positively or negatively act in provision of clinical benefit.

#### Conclusions

- **We have defined Proteomic maps of several batches of each of the Different IVIG formulations used in Australia**
  - **These reveal notable similarities and differences between preparations, including differences in the monomer to dimer Ig fractions and in the non-immunoglobulin proteins detected.**
    - **Direct identification of some of the different proteins present in the preparations, which were found to differ between preparations, was possible using mass spectrometry.**
      - **Cytokine multiplex-assay revealed additional variability in the cytokines detected in the respective IVIG formulations.**
        - **The important next step is to determine whether these differences are of therapeutic relevance, either as positive or negative effectors in IVIGs delivery of clinical benefit for a broad range of medical conditions.**

Acknowledgements: We wish to thank Dr Yasmin Ayob for IVIG. Reference: Gelfand et al Int Immunopharm 6: 592-599, 2006

Figure S1. Poster Presented at AHMR Congress 2006

*2005-2006 Conference presentations of these and closely related studies:*

***2005:*** G Lynch and J Sullivan. Application of Comparative Proteomics to the Study of Antibodies in IVIGs Prepared from Geographically Different Populations. 10th Annual *Australian Proteomics Symposium.* Cowes, Phillip Island, Victoria. Australia. 4-6th Feb.2005.

***2006:*** Lynch, G.W., Jeremy Kong, J., Wassinger, V., Bret Church, W.B., Sullivan, J., Michael Guilhaus, M. Applied Proteomic Studies for Comparison of Recombinant and Plasma-derived Therapeutic Proteins. 11th Annual *Australian Proteomics Symposium.* Lorne, Victoria. 3-5th Feb. 2006*. Australian Proteomics Symposium Proceedings.* Abstract 119.

Lynch, G.W, Fitzgerald A, Sullivan, J., Walsh, B. (2006) Proteomics in Transfusion Medicine and Virology: Characterisation of Intravenous Immunoglobulins and anti-Influenza Specificities. 11th Annual *Australian Proteomics Symposium. Lorne, Victoria. 3-5th Feb.2006.*

Lynch, G.W, Fitzgerald, A., Walsh, B., Kapitza, N., J. S. Sullivan, J.S. (2006) Translational Proteomics for Transfusion Medicine: Resolution of The IVIG Proteomes of Different Geographically Sourced and Prepared IVIG Immunotherapies. 26th Nov – 1st Dec. 2006. Melbourne, Victoria. *Australian Health and Medical Research Congress (AHMRCongress) Proceedings.* 5: 4.

Lynch, G.W., Kapitza, N., Fitzgerald, A., Walsh, B., Sullivan ,J. (2006) The anti-influenza specificities of intravenous immunoglobulins (IVIGs) for targeting human and avian influenza antigens: comparisons of northern and Southern Hemisphere IVIGs. 26th Nov – 1st Dec 2006. Melbourne, Victoria. *Australian Health and Medical Research Congress (AHMRCongress) Proceedings* 5: 467.

*Historic background of our Proteomic and Influenza-based studies of IVIGs, and wider implications:*

The studies described in this article were presented in poster format (Figure S1) at the AHMRCongress listed above, as were other related IVIG and Influenza directed antibody studies (Lynch et al 2006a), and other conference presentations of the findings elsewhere over the 2005–2006 period. The described IVIG studies, emerged initially from an early grounding on the clinical use and management of IVIG blood products by GL as a Senior Transfusion Medicine Scientist at the ARCBS in Sydney during the first half of 2004. This prompted a follow-up series of research studies by GL and JS to better understand and define the compositions, proteomic and biologic profiles of IVIGs from different Geographical regions, from mid 2004 and those interests continued to the present. In this endeavour, we have been ably assisted over the journey by many (> 30), student and technical collaborative projects along the way. Importantly the early studies and findings on IVIGs as described here, together with our Influenza studies, immediately revealed and exposed to us the existence of an otherwise hidden aspect of adaptive immunity. Further establishment of this alternate, but hidden new area of adaptive responses has been our significant challenge ever since and described in subsequent publications (e.g., Lynch et al 2008, Sullivan et al 2009, Lynch et al 2012). This has driven us to understand that the alternate mechanisms we have identified, if combined collectively together with traditionally well-known and described processes, could enable and provide a far more extensive adaptive realm of a governing *Immune Cosmos*©, than is generally considered. The molecular differences and mechanisms that drive these alternate forms of adaptive immunity indicated from our research have been elaborated further in reviews on the topic of broad *Seasoned Immunity*© (Lynch et al 2009, Lynch et al 2012). These are to our knowledge the first such reports, and support the contention, for the existence of two remarkably different dimensions of adaptive Immunity. We have termed them respectively, *Vivid Immunity*© and *Dark Immunity*©.

© GW Lynch, JS Sullivan: Theories of an overarching Immune Cosmos, comprising Vivid and Dark adaptive processes. Guided by separate demarcations of the: 1) *Theory of Vivid Adaptive Immunity'*© and 2) *Theory of Dark Adaptive Immunity'*©.

*Supplement References:*

Lynch, G.W., Kapitza, N., Fitzgerald, A., Walsh, B., Sullivan, J., (2006a) The anti-influenza specificities of intravenous immunoglobulins (IVIGs) for targeting human and avian influenza antigens: comparisons of northern and Southern Hemisphere IVIGs. Austr Health Med Res Congr Proc. 5: p. 467

Lynch, G.W., Selleck, P., Axell, A., Downton, T., Kapitza, N.M., Boehm, I., Dyer, W., Wang, Y., Stelzer-Braid, S., Rawlinson, W.D., Sullivan, J.S., (2008) Cross-Reactive anti-Avian H5N1 Influenza Neutralizing Antibodies in a Normal 'Exposure-Naïve' Australian Blood Donor Population. The Open Immunol J. 1(1). P. 13-19.

Sullivan, J.S., Selleck, P.W., Downton, T., Boehm, I., Axell, A-M., Ayob, Y., Kapitza, N.M., Dyer, W., Fitzgerald, A., Walsh, B., Lynch, G.W., (2009). Heterosubtypic anti-avian H5N1 influenza antibodies in intravenous immunoglobulins from globally separate populations protect against H5N1 infection in cell culture. J Mol Genet Med. 3(2): p. 217-24.

Lynch, G.W., Selleck, P., Church, W.B., Sullivan, J.S., (2012) Seasoned Adaptive Antibody Immunity for Highly Pathogenic Pandemic Influenza in Humans: Naturally acquired heterosubtypic humoral immunity can guide universal vaccines for novel influenzas. Immunonology and Cell Biology. 90: 149-158. doi:10.1038/icb.2011.38.

Lynch, G.W., P. Selleck, and J.S. Sullivan, *Acquired heterosubtypic antibodies in human immunity for avian H5N1 influenza.* (2009) J Mol Genet Med. 3(2): p. 205-9.

Lynch, G.W., Selleck, P., Sullivan, J.S., (2009) Acquired Antibody Immunity for Avian H5N1
Influenza. The Journal of Molecular and Genetic Medicine. 3 (2): 205-209. 2009
